# Comparative analysis of methodologies for detecting extrachromosomal circular DNA

**DOI:** 10.1101/2023.12.01.569546

**Authors:** Xuyuan Gao, Ke Liu, Songwen Luo, Meifang Tang, Nianping Liu, Chen Jiang, Jingwen Fang, Shouzhen Li, Yanbing Hou, Chuang Guo, Kun Qu

## Abstract

Extrachromosomal circular DNA (eccDNA) is crucial in oncogene amplification, gene transcription regulation, and intratumor heterogeneity. While various analysis pipelines and experimental methods have been developed for eccDNA identification, their detection efficiencies have not been systematically assessed. To address this, we evaluated the performance of 7 analysis pipelines using three simulated datasets, in terms of accuracy, similarity, duplication rate, and computational resource consumption. We also compared the eccDNA detection efficiency of 7 experimental methods through twenty-one real sequencing datasets. Our results identified Circle-Map and CReSIL as the most effective pipelines for eccDNA detection through short-read and long-read sequencing data, respectively. Moreover, third-generation sequencing-based Circle-Seq showed superior efficiency in detecting copy number-amplified eccDNA over 10 kb in length. These results offer valuable insights for researchers in choosing the most suitable methods for eccDNA research.

## Introduction

Sequencing-based studies have greatly advanced our understanding of extrachromosomal circular DNA (eccDNA), on its roles in oncogene amplification^1-4^, gene expression regulation^5^, genome rearrangements^6,7^, and intratumor heterogeneity^4^. Diverse analysis pipelines and experimental methods have been developed to detect eccDNA. Viraj Deshpande et al. introduced the AmpliconArchitect (AA) algorithm to predict amplicon structures and eccDNA from short-read (SR) whole-genome sequencing (WGS) (WGS-SR) data^8^. CReSIL utilizes coverage depths and breakpoint reads to identify eccDNA from long-read (LR) WGS (WGS-LR) data^9^. Kumar et al. developed Circle_finder to identify eccDNA from short-read ATAC-Seq (ATAC-Seq-SR) data by analyzing split reads for eccDNA coordinates^10^. However, the performance of these analysis pipelines might be limited by the data generated from the corresponding experimental methods. For example, WGS and ATAC-Seq may have low eccDNA detection efficiency because vast majority of the sequencing reads were generated from linear DNA, and WGS-SR can only detect the copy number amplified eccDNA (ecDNA)^4,6,11^.

To enhance eccDNA detection, researchers have developed methods such as Circle-Seq^7,12,13^ and 3SEP^14,15^ for eccDNA enrichment from crude DNA. Circle-Seq utilizes rolling circle amplification (RCA) for circular DNA amplification, whereas 3SEP employs Solution A for selective circular DNA recovery. Post-enrichment, eccDNA undergoes library construction for sequencing on platforms like Illumina (Circle-Seq-SR/3SEP-SR) or Oxford Nanopore Technology (ONT) (Circle-Seq-LR/3SEP-LR). Concurrently, various analysis pipelines have been developed to process eccDNA sequencing data. Circle-Map^16^, ECCsplorer^17^, Circle_finder^10^, and ecc_finder (map-sr)^18^ are tailored for short-read data analysis. For long-read data, pipelines such as CReSIL^9^, NanoCircle^7^, eccDNA_RCA_nanopore^14^ and ecc_finder map-ont mode are used. Additionally, ecc_finder offers de novo assembly options: Spades in the asm-sr mode and Tidehunter in the asm-ont mode as distinct algorithms to identify eccDNA from SR and LR sequencing profiles, respectively. These eccDNA-enriched methods and tailored pipelines facilitate eccDNA identification without reliance on copy number information^6^.

Choosing the most suitable analysis pipeline and experimental method for eccDNA research is a complex task. Existing evaluations of these pipelines often have limited scope, focusing on single aspects like accuracy^9^ or computational needs^18^, and rely on oversimplified simulations that fall short of representing the intricacies of actual sequencing data. Additionally, detection efficiency for specific eccDNA types varies significantly between enriched (such as Circle-Seq and 3SEP) and non-enriched experimental methods (such as WGS-SR, WGS-LR, and ATAC-Seq-SR). For example, the rolling circle amplification (RCA) step is known to preferentially amplify circular DNA under 10 kb^19^, while the bias of Solution A enrichment remains unclear.

To address these challenges, we conducted an in-depth evaluation of 7 analysis pipelines. The comparative analysis scopes included assessing accuracy (F1-score), similarity (PCC, RMSE, JSD), duplication rate, and computational resource cost using three simulated datasets designed to mirror real eccDNA characteristics. These datasets replicated the length distribution and chromosomal origins as previously identified^7,9^. Additionally, we compared the detection efficiencies of 7 methods on twenty-one real sequencing datasets for different eccDNA types. Our comparative analysis highlights the most effective pipelines for analyzing short-read and long-read data from eccDNA-enriched methods and underscores the variation in eccDNA detection efficiency across different experimental approaches. Our findings are intended to guide researchers in choosing the most suitable methodologies for their eccDNA studies and to foster the development of novel approaches for efficient eccDNA detection.

## Results

### Study design

To evaluate the performance of analysis pipelines in eccDNA identification, we developed a Python script to generate simulated eccDNA datasets. This script extrapolated length distribution, chromosomal origins, and chimeric eccDNA proportions from existing data to create a mix of simulated circular DNA (true positives) and linear DNA (true negatives). It also simulated the rolling circle amplification (RCA) process and subsequent sequencing on short-read (Illumina) and long-read (ONT) platforms (Figure 1A). Three simulated datasets were produced, mirroring eccDNA identified in human sperm cells^7^, EJM cell line^9^, and JJN3 cell line^9^ (Figure 1B and Supplementary Figures 1A, 1B), each comprising 10,000 circular and 10,000 linear DNA sequences at a depth of 50X.

**Figure 1.**
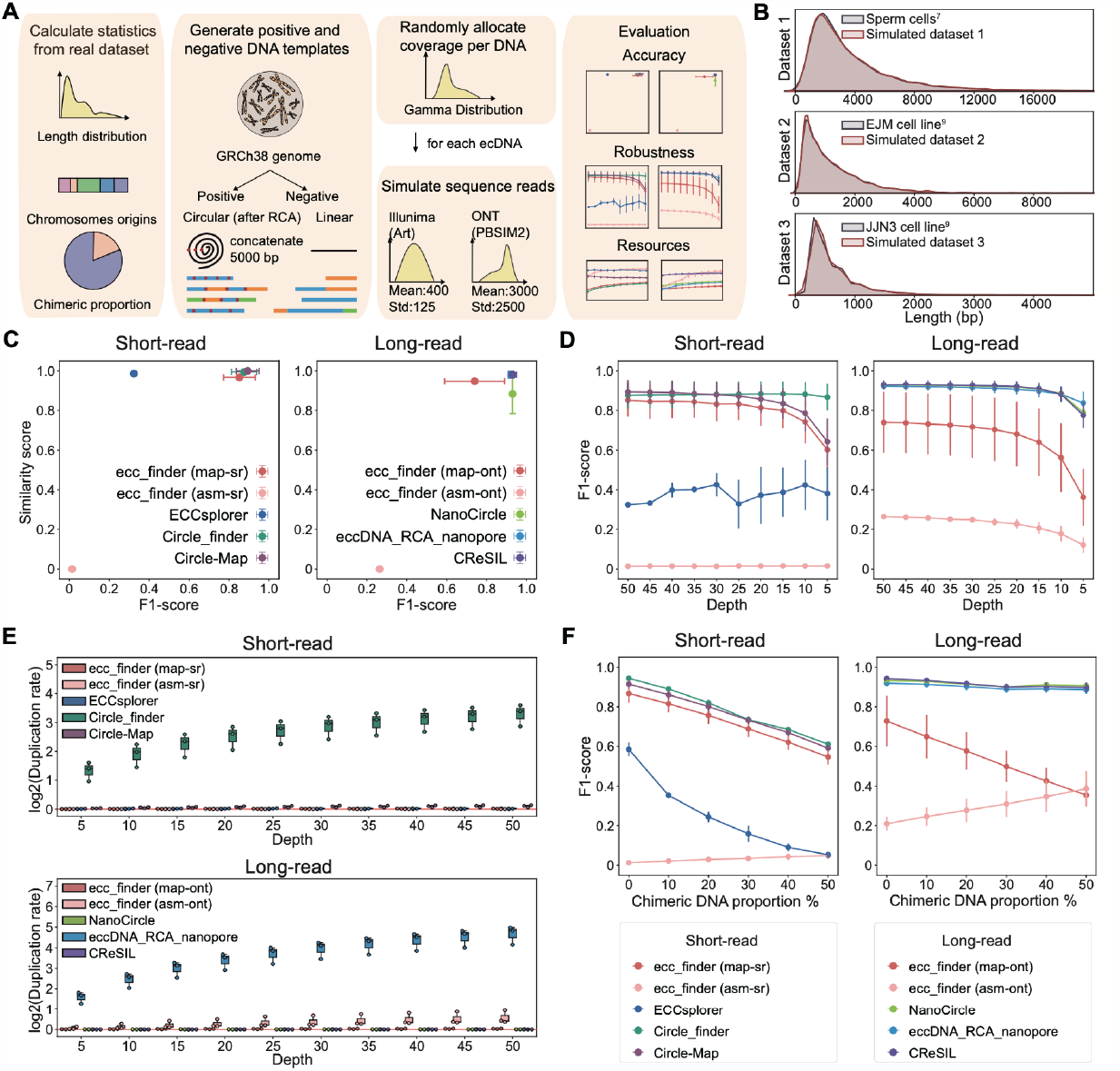
Assessment of analysis pipelines in eccDNA identification. **A**. Schematic overview of the benchmarking workflow used to compare the performance of bioinformatic pipelines. **B**. Length distribution comparison between simulated datasets and published datasets. **C**. Performance comparison of analysis pipelines at a simulated sequencing depth of 50X. **D**. Impact of simulated sequencing depth on eccDNA identification accuracy. **E**. Impact of simulated sequencing depth on duplication rates. **F**. Impact of chimeric DNA proportion on eccDNA identification accuracy.

We evaluated 10 modes of 7 pipelines, including Circle-Map, Circle_finder, ECCsplorer, and ecc_finder (map-sr/asm-sr) for short-read data, and CReSIL, eccDNA_RCA_nanopore, NanoCircle, and ecc_finder (map-ont/asm-ont) for long-read data. True positive identification was defined as over 90% sequence identity with the simulated eccDNA. Performance metrics included F1-score and a similarity score based on the root mean square error (RMSE), Pearson correlation coefficient (PCC), and Jensen–Shannon divergence (JSD) (see Methods). Additionally, we down-sampled the datasets to test pipeline robustness at low sequencing depths and generated datasets with varying chimeric DNA proportions (0-50%) to assess impact of chimeric DNA on eccDNA identification. We also introduced a duplication rate metric to address the issue of multiple detections of the same eccDNA sequence (see Methods) and analyzed the computational resource consumption for each pipeline.

For experimental method assessment, we selected Circle-Seq (SR/LR), 3SEP (SR/LR), WGS (SR/LR), and ATAC-Seq (SR) based on their non-targeted nature and sequencing compatibility with Illumina (SR) and ONT (LR) platforms (Figure 2A). To minimize batch effects, eccDNA was extracted from a uniform pool of HeLa cells. Controls included a pUC-19 plasmid (2686 bp) and a mouse *egfr* gene fragment (2651 bp), spiked into the cell lysate at a 1:1000 ratio to crude circular DNA. We then evaluated eccDNA detection efficiency of each method across various lengths and copy number statuses, quantifying detection efficiency as the number of eccDNA per gigabase (Gb) of sequencing data (see Methods).

**Figure 2.**
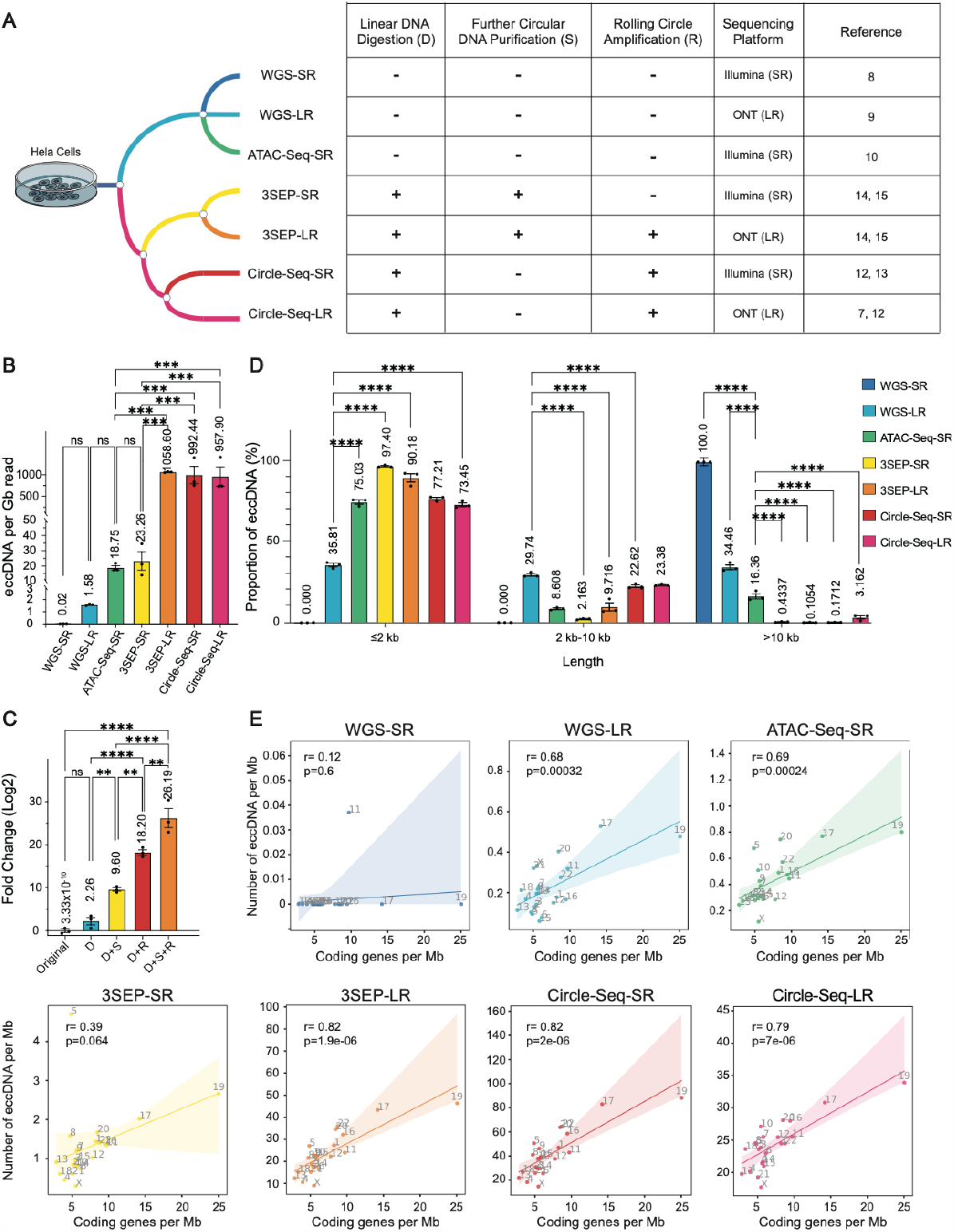
Impact of eccDNA enrichment operations on eccDNA identification. **A**. Schematic overview of the experimental methods comparison. **B**. eccDNA detection efficiency comparison. **C**. Ratio of spike-in plasmid DNA (pUC19) to linear DNA (*egfr* fragment) of the samples generated from different experimental steps. **D**. Detection efficiency for eccDNA with different length ranges. **E**. Correlation between eccDNA density and coding gene density. Dots represent individual experiments; n = 3 for all experiments. Statistical analyses were performed using one-way ANOVA (panel **B, C** and **D**, for panel **D** we also used one-way ANOVA analysis because we focused on the comparison within each length range), and Pearson correlation (panel **E**); error bars represent SEM. ∗p < 0.05, ∗∗p < 0.01, ∗∗∗p < 0.001, and ∗∗∗∗p < 0.0001.

### Assessment of analysis pipelines in eccDNA identification

In our evaluation of the performance of each analysis pipeline in eccDNA identification at a simulated sequencing depth of 50X, Circle-Map emerged as the most effective for short-read data, achieving an F1-score of 0.894 and a perfect similarity score of 1.00. Close contenders included Circle_finder, with an F1-score of 0.876 and similarity score of 0.994, and ecc_finder (map-sr), which scored an F1 of 0.852 and similarity of 0.967 (Figure 1C left panel). In the long-read data category, CReSIL led with an F1-score of 0.930 and a similarity score of 0.981, outperforming eccDNA_RCA_nanopore (F1-score: 0.923, Similarity Score: 0.980) and NanoCircle (F1-score: 0.929, Similarity Score: 0.884) (Figure 1C right panel). Besides, ecc_finder asm-sr mode was unable to identify eccDNA from SR data and the identified eccDNA from LR data by ecc_finder asm-ont mode had the lowest similarity score among all pipelines. Meanwhile, ECCsplorer could identify eccDNA from dataset 2 and 3 but failed in dataset 1 at sequencing depth 50X (Supplementary Table 1).

### Impact of sequencing depth on eccDNA identification

Previous research indicates that low eccDNA coverage adversely affects the performance of analysis pipelines in eccDNA identification^9^. To explore this, we down-sampled our three simulated datasets to various sequencing depths, assessing the performance of each pipeline in eccDNA identification. For short-read data, Circle-Map consistently achieved the highest F1-scores at sequencing depths above 30X (Figure 1D left panel). Below the depth of 30X, Circle_finder surpassed other pipelines in F1-score, and ECCsplorer could successfully identify eccDNA from dataset 1 (Figure 1D left panel & Supplementary Table 1). In the realm of long-read data, CReSIL led with the highest F1-scores at depths over 10X, while eccDNA_RCA_nanopore showed superior performance below a depth of 10X (Figure 1D right panel). Across all pipelines, the similarity score remained relatively stable, with minimal fluctuations (less than 0.2) as sequencing depth decreased from 50X to 5X (Supplementary Figure 1C). ecc_finder asm-sr mode could not identify the eccDNA across all the simulated sequencing depth (Figure 1D left panel), while asm-ont mode showed the lowest accuracy and similarity score among all the pipelines in analyzing LR data (Figure 1D right panel and Supplementary Table 1).

We observed a pattern of redundancy in eccDNA identification by eccDNA_RCA_Nanopore at all simulated depths, aligning with findings from another study^9^. Circle_finder also demonstrated similar redundancy. Upon calculating the duplication rates, it was evident that both Circle_finder and eccDNA_RCA_nanopore often identified multiple similar copies from a single eccDNA sequence (Figure 1E). These substantial duplication rates present considerable obstacles for the experimental validation of their predictions.

### Impact of chimeric DNA proportion on eccDNA identification

In addition to sequencing depth, we investigated the influence of chimeric DNA on eccDNA identification accuracy. We created simulated datasets with varying proportions of chimeric DNA, from 0% to 50%, maintaining a fixed sequencing depth of 20X. For short-read data, Circle-Map, Circle-finder and ecc_finder (map-sr) showed a similar decline in F1-score with increasing proportion of chimeric DNA, but were still the pipelines with top performance (Figure 1F left panel). ecc_finder (asm-sr) showed the lowest F1-score though it increased from 0.0128 at 0% chimeric DNA to 0.0487 at 50%. ECCsplorer experienced the most significant drop, with its F1-score falling from 0.585 at 0% to 0.053 at 50%. The similarity scores for these pipelines, however, remained relatively stable with fluctuations under 0.1 (Supplementary Figure 1D left panel). Among long-read data analysis pipelines, most maintained consistent F1-scores except for ecc_finder (map-ont) and ecc_finder (asm-ont) (Figure 1F right panel). Their similarity scores also exhibited stability, except for NanoCircle, which showed greater fluctuation (Supplementary Figure 1D right panel).

These results demonstrate that an increased proportion of chimeric DNA can impact the accuracy of eccDNA identification by analysis pipelines. Delving further, we evaluated the recall rates for simple and chimeric eccDNA across pipelines. Circle-Map, Circle_finder, ECCsplorer, ecc_finder (map-sr/map-ont) had zero recall for chimeric eccDNA, suggesting their inability to detect chimeric eccDNA (Supplementary Figure 1E). Specifically, ECCsplorer’s recall for simple eccDNA plummeted from 0.420 to 0.054 as the proportion of chimeric DNA rose from 0% to 50%. Besides, ecc_finder (asm-sr/asm-ont) exhibited higher recall rates for chimeric eccDNA compared to simple eccDNA (Supplementary Figure 1E).

### Computational resources consumed by different analysis pipelines

In our evaluation of computational resources consumed by each pipeline, we utilized a computer cluster equipped with two Intel Xeon Scale 6248 CPUs (2.5 GHz, 320 CPU cores), 384 GB of DDR4 memory, and 2 TB AEP memory. We observed that both the time and memory consumption of most pipelines increased with mean coverage rising (Supplementary Figures 1F & 1G). Notably, Circle-Map and CReSIL kept stable computational resources consumption across all sequencing depths (Supplementary Figure 1F & 1G). ECCsplorer experienced memory errors on our platform when analyzing dataset 1 (eccDNA from human sperm cells) at sequencing depths above 25X (Supplementary Figure 1F right panel).

Based on the above analysis, we concluded that Circle-Map and CReSIL were the most appropriate analysis pipelines to analyze eccDNA-enriched short-read and long-read data, respectively, due to their high detection accuracy, high similarity score, low duplication rate and stable computational resource consumption. We thereby applied them to benchmark the 7 experimental methods for their efficiency of eccDNA identification (Figure 2A).

### Impact of eccDNA enrichment steps on eccDNA identification

We assessed eccDNA detection efficiency by the number of eccDNA detected per gigabyte (Gb) of data. The results indicated that methods incorporating RCA steps achieved significantly higher eccDNA detection efficiencies compared to those without RCA (Figure 2B). Notably, qPCR analyses revealed that both Solution A purification and the RCA step considerably increased the log2 ratio of circular to linear spike-in DNA (Solution A: from 2.26 to 9.60 and from 18.20 to 26.19, RCA: from 2.26 to 18.20 and from 9.60 to 26.19) (Figure 2C). To validate these findings, we randomly selected nine simple and seven chimeric eccDNA for testing, observing validation rates above 0.6 in RCA-utilizing methods (3SEP-LR: 10/16, Circle-Seq-SR: 8/9, Circle-Seq-LR: 11/16) (Supplementary Figure 2 & Supplementary Table 2).

Further analysis of the eccDNA length distribution and chromatin origins revealed that Circle-Seq-LR had the highest detection efficiency for >10 kb eccDNA and enriched methods (except for 3SEP-SR) could detect significantly more <=10 kb eccDNA per Gb data than non-enriched methods (Figure 2D). However, over 97% of the identified eccDNA from eccDNA-enriched methods were shorter than 10 kb (Circle-Seq-LR: 97%, Circle-Seq-SR: 99.8%, 3SEP-LR: 99.9%, 3SEP-SR: 99.5%) and over 90% of eccDNA detected by methods like 3SEP-SR and 3SEP-LR were shorter than 2 kb (Supplementary Figure 3). In contrast, non-enriched methods showed a higher proportion of eccDNA lengths exceeding 10 kb (Supplementary Figure 3). Additionally, except for 3SEP-SR and WGS-SR, a significant positive correlation was observed between eccDNA density (number of detected eccDNA per million base (Mb)) and protein-coding gene density across chromosomes in most methods, consistent with prior studies^7,13^ (Figure 2E). 3SEP-SR showed a similar trend, though the correlation was not statistically significant (r=0.39, p=0.064), and no significant correlation was found in WGS-SR data (r=0.12, p=0.6). This could be due to the limited number of eccDNA identified by WGS-SR, suggesting the importance of eccDNA enrichment in experimental setups to obtain a comprehensive eccDNA profile.

### Detection efficiency of ecDNA by different experimental methods

The eccDNA overlapping with copy number amplified regions was designated as ecDNA, while eccDNA outside these regions was categorized as nonecDNA^11^. Circle-Seq-SR, Circle-Seq-LR, and 3SEP-LR identified a higher average number of ecDNA per Gb of data (205.2, 165.8, and 203.9, respectively) compared to WGS-SR, WGS-LR, and ATAC-Seq-SR (0.01576, 0.9100, and 6.862, respectively) (Figure 3A). However, a significantly higher proportions of ecDNA were found in the eccDNA detected by WGS-SR (100%), WGS-LR (57.68%), and ATAC-Seq-SR (36.67%) compared to Circle-Seq-SR (20.58%), Circle-Seq-LR (17.09%), and 3SEP-LR (19.26%) (Figure 3B).

**Figure 3.**
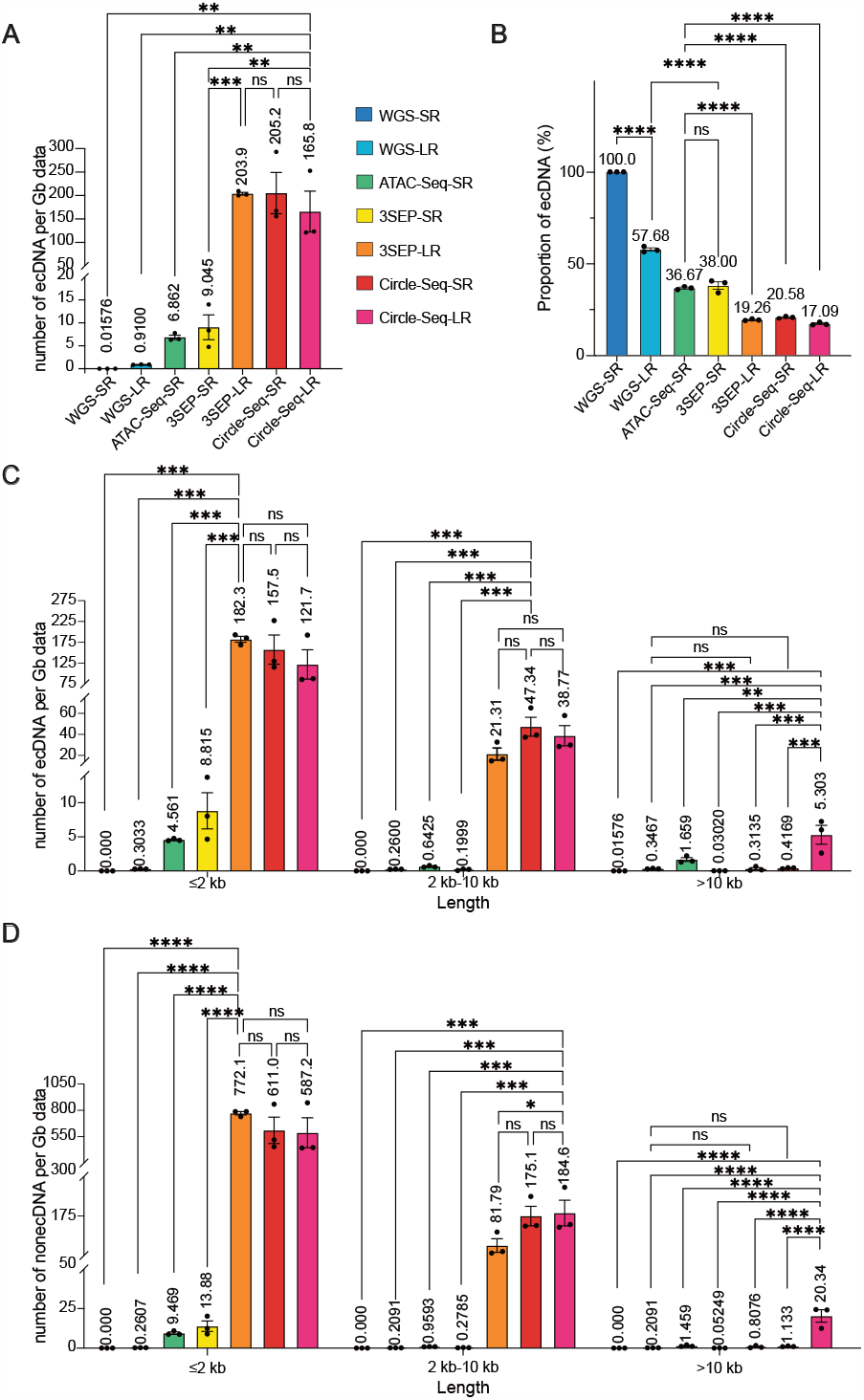
Detection efficiency of ecDNA by 7 experimental methods. **A**. ecDNA detection efficiency of 7 experimental methods. **B**. Comparison of the proportion of ecDNA in the total detected eccDNA. **C**. Comparison of the detection efficiency of ecDNA with different length ranges by 7 experimental methods. **D**. Comparison of the detection efficiency of nonecDNA with different length ranges by 7 experimental methods. Dots represent individual experiments; n = 3 for all experiments. Statistical analyses were performed using one-way ANOVA (for **C** and **D** we used one-way ANOVA analysis because we focused on the comparison within each length range); error bars represent SEM. ∗p < 0.05, ∗∗p < 0.01, ∗∗∗p < 0.001, and ∗∗∗∗p < 0.0001.

Subsequently, we further analyzed the detection efficiencies for both ecDNA and nonecDNA across varying lengths (<=2 kb, 2-10 kb, >10 kb Figure 3C & 3D). 3SEP-LR demonstrated the highest efficiency in detecting both ecDNA and nonecDNA up to 2 kb in length. Circle-Seq-SR was the most efficient for detecting ecDNA between 2 kb and 10 kb. For eccDNA over 10 kb, Circle-Seq-LR outperformed all other methods in detecting both ecDNA and nonecDNA. Interestingly, for detecting ecDNA and nonecDNA over 10 kb, WGS-LR, despite not employing a circular DNA enrichment step, showed comparable efficiency with 3SEP-SR, 3SEP-LR, and Circle-Seq-SR (Figures 3C & 3D).

## Discussion

Benchmarking the available analysis pipelines and experimental protocols for detecting eccDNA is crucial for advancing eccDNA research. In this study, we have identified key performers for eccDNA detection by assessing 7 analysis pipelines using various metrics, and comparing 7 experimental methods via detection efficiency. Circle-Map and CReSIL stand out for their ability to identify eccDNA from short-read and long-read data, respectively. In the realm of experimental methods, Circle-Seq-LR demonstrates the highest detection efficiency for longer eccDNA, while 3SEP-LR is more effective for shorter eccDNA. This information is vital for researchers in selecting the most suitable methodologies for their eccDNA studies.

Despite our simulated datasets closely mimicked the length distribution of real eccDNA data, they featured a comparatively smaller proportion of eccDNA longer than 10 kb. This imbalance posed challenges in precisely evaluating the performance of different analysis pipelines across various eccDNA length ranges. Additionally, while using DNA from a cell line sheds light on the eccDNA detection efficiency of diverse methods, the potential copy number bias introduced at different experimental stages remains a concern due to the absence of a known ground truth. Future research could benefit from employing a specially designed circular DNA pool with a defined copy number. Such a controlled approach would not only help in addressing potential biases but also allow for more accurate quantification of metrics like F1-score and similarity score for each experimental method in eccDNA detection.

Split and discordant reads within short-read data, and breakpoint reads in long-read data, are primary sources for eccDNA identification. CReSIL utilizes the breakpoint read information to construct directed graphs, allowing for its effective identification of eccDNA from both the concatemeric tandem copies (CTC) reads and the non-CTC reads containing breakpoints. Conversely, eccDNA_RCA_nanopore only focuses on CTC reads and might limit its ability to identify larger eccDNA that were hard to generate CTC reads. Both eccDNA_RCA_nanopore and Circle_finder exhibit a tendency for redundancy due to their approach of reporting results for each CTC read or split read, respectively. Our study also shows that ecc_finder is uniquely capable of identifying chimeric eccDNA from short-read data, owing to its asm-sr mode, but the asm-sr and asm-ont modes of the same tool might not be suitable for identifying eccDNA from eccDNA-enriched data. Because the available pipelines are limited for analyzing eccDNA non-enriched data, we only compared the performance of these analysis pipelines for identifying eccDNA from simulated eccDNA-enriched datasets. Future study is needed to compare the performance of the analysis pipelines for detecting eccDNA from non-enriched data when more pipelines are available.

This benchmark study also helps to explain controversial findings in the field. For instance, the limited detection of ecDNA in normal cells^4^ may be due to the low sensitivity of WGS-SR in identifying eccDNA. Conversely, the effective identification of eccDNA in human germline cells may be facilitated by the use of the Circle-Seq-LR technique^7^. However, it is important to note from our analysis that non-enriched methods like WGS-SR hold their own unique advantages, such as providing copy number variation information essential for ecDNA classification^20^. Therefore, we do not suggest that non-enriched methods be replaced by enriched methods. Moreover, other non-enriched methods like WGS-LR^21^ and modified ATAC-Seq-SR^22^ can preserve nucleotide decorations in the sequencing reads, a feature could potentially lost in sequences generated from enrichment steps like RCA.

A significant challenge in eccDNA research is the inconsistency in the definitions of different eccDNA types used by various studies. We defined ecDNA as eccDNA colocalizing with genome copy number-amplified regions^11^, due to the putative gene amplification effect of ecDNA. Other studies may use size thresholds to define ecDNA^23,24^. Establishing a consensus definition is crucial for harmonizing research findings in this rapidly evolving field.

Lastly, the potential of eccDNA as a diagnostic marker for diseases like advanced chronic kidney disease^25^, medulloblastoma^26^, and colorectal cancer^27^ is promising. However, the time-consuming enrichment step, particularly the linear DNA digestion process, may limit the practicality of eccDNA in clinical diagnostics. Our findings suggest that linear DNA digestion has a marginal effect compared to RCA or Solution A purification, and future method developments might consider omitting this step to reduce time costs. Alternatively, skipping linear DNA digestion and purifying circular DNA and RCA products using Solution A could be explored, though this might preferentially enrich shorter eccDNA. Optimizing the RCA step, typically a lengthy process, could also enhance the feasibility of using eccDNA information for clinical diagnosis.

## Methods

### Generation of simulated datasets

To generate simulated eccDNA datasets for evaluation, we created a python script to simulate datasets, containing circular and linear DNA, according to the length distribution, chromosome origins and chimeric eccDNA proportion of the eccDNA from the given data. We collected the eccDNA identified from human sperm cells^7^, EJM cell line^9^, and JJN3^9^ cell line and used these three datasets as input. We generated three simulated datasets, containing 10000 circular DNA (as positive sequences) and 10000 linear DNA fragments (as negative sequences). Then, we randomly shifted the positive sequence to mimic the breakpoint of eccDNA and concatenated the 5000 bp of individual simulated eccDNA to mimic the RCA procedure. We used generated sequences as templates to further simulate short-read datasets using ART^28^ (--sr-platform ‘HS25’ --sr-mean ‘400’ --sr-std ‘125’ --sr-readlen ‘150’) and simulate long-read datasets using PBSIM2^29^ (--ont-model ‘R94’, --ont-mean ‘3000’,--ont-std ‘2500’) with different sequencing depth (5X, 10X, 15X, 20X, 25X, 30X, 35X, 40X, 45X, 50X). We also simulated short-read datasets and long-read datasets with different chimeric DNA ratios (0%, 10%, 20%, 30%, 40%, 50%) at sequencing depth 20X.

### Performance evaluation of each pipeline

The identification of eccDNA was done following the instructions on the website of each pipeline. We used hg38 genome as reference. For Circle-Map^16^, we used Circle Map Realign to identify eccDNA and used recommended filters (circle score > 50, split reads > 2, discordant reads > 2, coverage increase in the start coordinate > 0.33 and coverage increase in the end coordinate > 0.33). For Circle_finder^10^, we used the script circle_finder-pipeline-bwa-mem-samblaster.sh to identify eccDNA. For ECCsplorer^17^, we used mapping module to identify eccDNA. For ecc_finder^18^, all the 4 modes were used to identify eccDNA from either short-read or long-read data. The identified eccDNA with length longer than 10^7^ bp was filtered out. For CReSIL^9^, we followed the instruction on its website to identify eccDNA and considered cyclic eccDNA as identified results. For NanoCircle^7^, we followed the instruction on its website and considered high_conf simple eccDNA and complex eccDNA as identified results. For eccDNA_RCA_nanopore^14^, we followed the instruction on its website to identify eccDNA. For the pipelines that did not supply FASTA format results, we used pysam^30^ to transform bed format into FASTA format. The FASTA files were then compared to the simulated eccDNA sequence by MUMmer3^31^.

### Cell culture

HeLa cells were bought from BeNa Culture Collection (Cat#BNCC342189; RRID: CVCL-0030). NIH3T3 (RRID: CRL-1658) was a gift from Prof. Shu Zhu lab of the University of Science and Technology of China. HeLa cells or NIH3T3 cells were cultured at 37°C in DMEM (Thermo Fisher Scientific 11965092) containing 10% FBS (Thermo Fisher Scientific 10091148) and 1% penicillin–streptomycin (Thermo Fisher Scientific 15140122). Upon reaching approximately 80%-100% confluence, the cells were rinsed with 1× PBS (Sangon Biotech, B540626-0500) and digested with 0.25% trypsin (Beyotime C0203-500 ml). The trypsinization process was terminated by adding DMEM+10% FBS+1% penicillin– streptomycin, and the cells were collected by centrifugation at 500×g for 5 min at RT. Cells were then washed twice by using 1X PBS and then centrifuged at 500xg for 5 min at 4°C to obtain the cell pellet for following experiments.

### ATAC-seq library construction

For each replicate, approximately 50000 cells and a commercialized Tn5 kit (Vazyme, TD501) were used to construct the ATAC-Seq library. The reaction mix, consisting of 50,000 cells, 0.005% digitonin (Sigma–Aldrich D141-100MG), 33 mM Tris-Ac (pH 7.8), 66 mM KAc, 10 mM MgAc, and 16% DMF, was incubated at 500 rpm for 30 mins at 37°C using a thermal rotator. After the reaction, the cells were washed twice using wash buffer (10 mM Tris-HCl pH 7.5, 10 mM NaCl, 3 mM MgCl_2_, 0.005% digitonin) and resuspended in 14 µl of 10 mM Tris-HCl pH 7.5. Cells were then lysed by mixing with 2 µl lysis buffer (200 mM Tris-HCl pH 8.0, 0.4% SDS) and 0.2 µl proteinase K (20 mg/mL) at 500 rpm for 15 mins at 55 °C. The lysis reaction was terminated by adding 4 µL of 10% Tween-20 and 0.4 µL of 100 mM PMSF. The samples were incubated for 5 mins at RT, and then PCR was performed to add adapters to the DNA segment for sequencing.

### Whole-genome sequencing

For preparing each replicate for WGS-SR, after washing the cells, more than 1 million cells were frozen using liquid nitrogen. Three replicates were sent to Sequanta Technologies for library construction and WGS-SR sequencing (Illumina NovaSeq 6000 platform). For preparing each replicate for WGS-LR, after washing the cells, more than 5 million cells were frozen using liquid nitrogen. Three replicates were sent to Novogene for library construction and WGS-LR sequencing (Oxford Nanopore PromethlON platform).

### Isolation of crude circular DNA

Crude circular DNA was extracted from the same pool of HeLa cells. The details were described in the published protocol^15^. In brief, more than 60 million HeLa cells were used to extract the crude circular DNA pool. For each reaction (approximately 30 million HeLa cells), cells were collected in a 50 mL tube by centrifugation at 2,000xg for 10 mins at 4°C. Resuspend the cells in 10 ml of suspension buffer (10 mM EDTA pH8.0, 150 mM NaCl, 1% glycerol, Lysis blue (1×, from QIAGEN Plasmid Plus Midi Kit), RNase A (0.55 mg/ml), and freshly supplemented with 20 µL of 2-mercaptoethanol). Add 10 mL Pyr buffer (0.5M pyrrolidine, 20 mM EDTA, 1% SDS, adjust pH to 11.80 with 2 M Sodium Acetate pH 4.00, and freshly supplemented with 20 µL 2-mercaptoethanol) to the cell suspension. Gently mix by inverting the tube 5-10 times and incubate at room temperature for 5 mins. After lysis, 10 mL of Buffer S3 (From QIAGEN Plasmid Plus Midi Kit) was added to the mixture, and the tube was gently inverted until the solution color turned white. Then, the lysate was centrifuged at 4500xg for 10 mins. The clear lysate was transferred to a QIAilter Catridge (From QIAGEN Plasmid Plus Midi Kit) and incubated at room temperature for 10 mins. Then, the cell lysate was filtered into a 50 mL tube. The volume of the filtrated lysate was approximately 27 mL, and 9-10 mL of Buffer BB (1/3 of the lysate volume, From QIAGEN Plasmid Plus Midi Kit) was added. The lysate was mixed by inverting the tube 4-8 times. The lysate mixture was then transferred to the spin column, and vacuum was applied until all liquid passed through. We added 0.7 mL ETR buffer (From QIAGEN Plasmid Plus Midi Kit) to wash the column, and applied vacuum until all liquid passed through. Then, the wash was repeated by using 0.7 mL PE buffer (From QIAGEN Plasmid Plus Midi Kit). After washing, the tube was centrifuged at 10000xg for 2 mins to remove the liquid, and the column was transferred to a new clean 1.5 mL centrifuge tube. Crude eccDNA was then eluted by using 100 µL of 0.1x EB buffer (From QIAGEN Plasmid Plus Midi Kit). For each microgram crude eccDNA we spiked in 1 ng pUC19^32^ (was a gift from Joachim Messing, Addgene plasmid # 50005; RRID: Addgene_50005) and 1 ng *egfr* fragment to generate crude circular DNA mixture.

### Linear DNA digestion

For each DNA digestion reaction, 3 µg crude circular DNA mixture was digested by using 0.5 µL Pac I and 1 µL ATP-dependent Plasmid Safe DNase in 1X ATP-dependent Plasmid-Safe DNase buffer. Then, 0.1 µL of 110 mg/ml RNase A and 2 µL of 25 mM ATP were added to the reaction in a total volume of 50 µL. The reaction mix was incubated at 37°C for 16 hours. After digestion, 1.8X SPRIselect beads were used to purify the DNA. DNA was eluted with 66 µL of 2 mM Tris-HCl pH=7.0 to carry out Solution A purification or eluted with 66 µL of 0.1 X EB buffer (From QIAGEN Plasmid Plus Midi Kit) without further Solution A purification.

### Solution A purification

The Solution A purification step followed the published study^15^ and was used in 3SEP-SR and 3SEP-LR only. In brief, we transferred 50 µL eluted circular DNA (in 2mM Tris-HCl pH=7.0) to a 1.5 mL tube. Added 700 µL of Solution A (room temperature) to the tube, mixed by pipetting up and down, and incubated at room temperature for 5 mins. Took 10 µL Dynabeads™ MyOne™ Silane beads (resuspend by thoroughly vortex) to a 200 µL tube and stood it on a magnetic shelf. When beads were settled, removed the liquid and added 20 µL Solution A to resuspend the beads. Then we transferred the beads to DNA (incubated in Solution A) and pipetted up and down for 10 times. Put the mixture on a magnetic shelf, and removed the liquid when the beads were settled. Quickly spun down the beads and put it on the magnetic shelf again to remove the residual liquid. Took off the tube from magnetic shelf and resuspended the beads in 300 µL Solution A. Put the tube on the magnetic shelf and removed the liquid when the beads were settled. Quickly spun down the beads and put it on the magnetic shelf, removing the residual Solution A when beads were settled. Repeated the 300 µL Solution A wash once more. After the second Solution A wash, kept the tube on the magnetic shelf, added 700µL 3.5M NaCl, waited for 1 minute and then removed the liquid, and repeated once. After the second NaCl wash, kept the tube on the magnetic shelf, added 800µL freshly prepared 80% ethanol, waited for 1 minute and then removed the liquid, and repeated once. Quickly spun down the beads and put it on the magnetic shelf again to remove the residual liquid. Took off the tube and used 30 µL 0.1X EB buffer (From QIAGEN Plasmid Plus Midi Kit) to resuspend the beads and incubated for more than 3 minutes. Put the tube back to the magnetic shelf and transferred the elute (contained purified circular DNA) when beads were settled.

### Rolling Cycle Amplification (RCA) and debranching

We measured the DNA product concentration by using Qubit 4.0, and aliquoted 1 ng DNA to prepare the RCA reaction premix (2 µL 10X Phi 29 DNA Polymerase Reaction Buffer, 2 µL dNTPs (25 mM each), 1 µL Exo-resistant Random Primer, and add H_2_O to 17.6 µL). The samples were incubated at 95°C for 5 mins and then ramped to 30°C at -0.1°C/sec. Then, added 1 µL of Phi29 DNA Polymerase, 1 µL of Pyrophosphatase (Inorganic) and 0.4 µL of recombinant Albumin (offered with Phi 29 DNA polymerase) to a 20 µL final reaction mix. The samples were incubated at 30°C for 14 hours and inactivated at 65°C for 10 mins. The product was diluted by adding 80 µL of H_2_O, and 1.8X SPRIselect beads were used to purify the product. Eluted the DNA product in 0.1X EB (From QIAGEN Plasmid Plus Midi Kit) buffer. T7 endonuclease I was employed to cleave the branched RCA product from circular DNA. Briefly, 6 µg RCA product was aliquoted into the reaction tube along with 30 µL 10X NEBuffer 2 and 15 µL T7 Endonuclease I, and H2O was added to 300 µL. The reaction mix was incubated at 37°C for 15 mins. Used 0.4X SPRIselect to purify the reaction product.

### DNA fragmentation

For Circle-Seq-SR, the debranched DNA materials were sent to Sequanta Technologies for ultrasonic fragmentation with the fragment size in 300-500 bp as reported in the published protocol^12^. For 3SEP-SR, the Solution A purified DNA material was sent to Sequanta Technologies for enzymatic fragmentation. To compare across different experimental methods, 1 ng DNA was used to generate the sequencing library by using Nextera XT DNA Library Preparation Kit (Illumina).

### Sequencing

For ATAC-Seq-SR, 3SEP-SR, and Circle-Seq-SR, DNA library was sequenced by Sequanta Technologies on Illumina NovaSeq 6000 platform. For 3SEP-LR and Circle-Seq-LR, the long-read sequencing library was constructed by Novogene and sequenced on Oxford Nanopore PromethlON platform.

### Identification of eccDNA from real datasets

We used the script circle_finder-pipeline-bwa-mem-samblaster.sh in Circle_finder^10^ to identify eccDNA from ATAC-seq-SR data and set a filter (length shorter than 10^7^ bp) to select eccDNA. For WGS-SR data, we used AmpliconArchitect^8^ to identified eccDNA with options (cngain=4, cnsize=10000). For WGS-LR data, we used CReSIL identify_wgls command^9^ to identify eccDNA, and filtered cyclic eccDNA. For Circle-seq-SR and 3SEP-SR data, we used Circle Map Realign^16^ to identify eccDNA and used recommended filters (circle score > 50, split reads > 2, discordant reads > 2, coverage increase in the start coordinate > 0.33 and coverage increase in the end coordinate > 0.33, length<10^7^bp). For Circle-seq-LR and 3SEP-LR data, we used CReSIL identify command^9^ to identify eccDNA and filtered cyclic eccDNA.

### Identification of ecDNA

We used Control-FREEC^33^ (breakPointThreshold = 0.6, window = 50000, step=10000) to examine the copy number variation in 3 replicates of our WGS-LR data. We defined eccDNA as ecDNA if it had overlap with the CNV gain regions identified by Control-FREEC.

### PCR validation

DNA sequences spanning the breakpoint were obtained by using Genome Browser (https://genome.ucsc.edu/index.html). Primers targeting the eccDNA breakpoint were designed by using Primer-Blast (https://www.ncbi.nlm.nih.gov/tools/primer-blast/) (Supplementary Table 2). The Hela cell genome was extracted by using the DNeasy^®^ Blood & Tissue Kit (QIAGEN Cat. No. 69504). KOD FX (TOYOBO No. KFX-101) was used to perform the PCR. In brief, 20 ng DNA template (Genome DNA or Sample), 1.5 µL 10 µM forward primer, 1.5 µL 10 µM reverse primer, 4 µl 2 mM dNTPs, 10 µL 2X PCR Buffer for KOD FX, 1 µL KOD FX and nuclease-free water (Invitrogen 10977015) (to a 20 µL final volume) were combined. PCR was carried out by using the following thermal cycle: 94°C for 2 minutes and then 30 cycles at 98°C for 10 s, 60°C for 30 s, 68°C for 1 minute and 68°C for 5 minutes. The PCR product was cut from the electrophoresis gel and sent for Sanger sequencing validation (by Sangon Biotech).

### Benchmark metrics

#### 1. F1-score

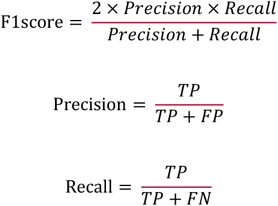

Then, we used the Pearson correlation coefficient (PCC), root mean square error (RMSE) and Jensen–Shannon divergence (JSD) of the length of the intersection and the union of the identified region and simulated region to evaluate the similarity of the identified eccDNA. The calculation of Jensen–Shannon divergence (JSD) is based on relative information entropy (that is, Kullback–Leibler divergence (KL)). The higher the PCC is, the more accurate the identified region. Lower RMSE and JSD values represent higher accuracy. We defined a similarity score by aggregating the PCC, RMSE and JSD. The pipeline with the best performance in each metric would have a value of 1, and the pipeline with the worst performance would have a value of 0. The score of other pipelines is determined by linear integration.

#### 2. PCC

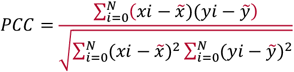

where *N* is the number of identified eccDNAs, *xi* is the length of the intersection of the identified and simulated regions and *yi* is the length of the union of the identified and simulated regions.

#### 3. RMSE

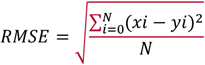

where *N* is the number of identified eccDNAs, *xi* is the length of the intersection of the identified and simulated regions and *yi* is the length of the union of the identified and simulated regions.

#### 4. JSD

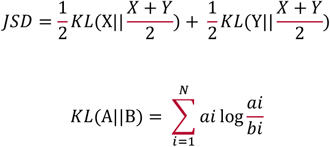

#### 5. Similarity Score

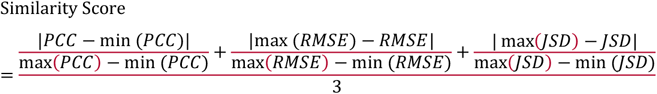

#### 6. Duplication Rate

The duplication rate is defined by the number of identified eccDNA (TP2) that have at least a 90% overlap of simulated eccDNA divided by the number of simulated eccDNAs (TP1) that can be identified by each pipeline.

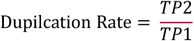

#### 7. Detection efficiency of specific type of eccDNA

Detection efficiency of specific type of eccDNA (per Gb) was calculated by using the following formula:

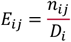

Where:

*E*_*ij*_: detection efficiency of experimental method *i* in detecting eccDNA type *j*

*n*_*ij*_: number of eccDNA in type *j* detected by experimental method *i*

*D*_*i*_: Size of the data (Gb) generated by experimental method *i*

### Statistics & Reproducibility

For performance evaluation of bioinformatic pipelines. We used Seaborn^34^ to visualize statistical data. Each point showed the Mean ± SEM (Standard Error of the Mean) in the figure. For column chart, one-way ANOVA (by GraphPad Prism 9) was used to evaluate the statistical significance (degrees of freedom between methods are 6, and degrees of freedom within methods are 14). For group column chart we also used one-way ANOVA (degrees of freedom between methods are 6 and degrees of freedom within methods are 14), because we focused on the comparison within each length range. Each column showed the Mean ± SEM and data points were shown as black dot on the column. Significant P values were indicated as follows: P ≤0.05 (∗), P ≤0.01(∗∗) and P ≤0.001(∗∗∗), P≤0.0001(∗∗∗∗). For correlation dot plot (Figure 2e), we used Pearson correlation in scipy.stats^35^ to measure the linear relationship between the density of coding genes and the density of eccDNA for each chromosome, and used Seaborn to present the result.

## Data Availability

The raw sequence data (WGS-SR, WGS-LR, ATAC-Seq-SR, 3SEP-SR, 3SEP-LR, Circle-Seq-SR, and Circle-Seq-LR) reported in this paper have been deposited in the Genome Sequence Archive^36^ in National Genomics Data Center^37^, China National Center for Bioinformation / Beijing Institute of Genomics, Chinese Academy of Sciences (GSA-Human: HRA006020) that are publicly accessible at https://ngdc.cncb.ac.cn/gsa-human. Any additional information required to reanalyze the data reported in this paper is available from the corresponding author upon request (Kun Qu).

## Code Availability

All original code has been deposited at Github (https://github.com/QuKunLab/eccDNABenchmarking). The simulated datasets can be generated by using the uploaded code. Any additional information required to reanalyze the data reported in this paper is available from the corresponding author upon request (Kun Qu).

## Acknowledgments

We thank all members in the Qu laboratory for helpful discussions. We are grateful for the gift of NIH3T3 cells from Prof. Shu Zhu of the University of Science and Technology of China. This work was supported by the National Key R&D Program of China (2020YFA0112200 and 2022YFA1303200 to K.Q.), the National Natural Science Foundation of China grants (T2125012 and 91940306 to K.Q.), CAS Project for Young Scientists in Basic Research YSBR-005 (to K.Q.), Anhui Province Science and Technology Key Program (202003a07020021 to K.Q.) and the Fundamental Research Funds for the Central Universities (YD9100002032 and WK9110000141 to K.Q.). We thank the USTC Supercomputing Center and the School of Life Science Bioinformatics Center for providing computing resources for this project.

## Author contributions

K.Q. and J. F. conceived the project. X.G., K.L., S.L., S.Z.L. and J.F. designed the framework. X.G. and S.L. performed all the wet-lab experiments with helps from M.T, S.Z.L., Y.H. and C.G.; K.L. and S.L. performed all the bioinformatics analysis with helps from N.L. and C.J.; X.G., K.Q., K.L., and S.L. wrote the manuscript with inputs from all authors. K.Q. supervised the project.

## Declaration of generative AI and AI-assisted technologies in the writing process

During the preparation of this work the authors used ChatGPT 3.5 and ChatGPT 4.0 in order to improve the language and readability. After using these tools, the authors reviewed and edited the content as needed and take full responsibility for the content of the publication.

